# Effect of recombinant phycocyanin on photosynthetic system in *Chlamydomonas reinhardtii*

**DOI:** 10.1101/813816

**Authors:** Feng Zhang, Xiaomei Cong, Zhendong Wang, Yalin Guo, Lulu Hou, Rui Li, Menghui Shang, Xuehong Wei, Zhaxi Yangzong, Xiaoting Xu, Xiaonan Zang

**Author notes:** Corresponding author: Xiaonan Zang.

## Abstract

The phycobilisome is an important photosynthetic antenna in the photosynthetic cyanobacteria, and phycocyanin is one of the main components of phycobilisomes. It helps cells absorb green light that green-lineage photo-synthetic organisms cannot. In this work, phycocyanin, heme oxidase and ferredoxin oxidoreductase from *Arthrospira platensis* FACHB 314 were successfully expressed in the green algae *Chlamydomonas reinhardtii*. Then the effects of this expression on the photosynthesis and growth of *C. reinhardtii* were detected. Transcriptional level analysis showed that the phycocyanin gene was successfully expressed stably in the transgenic strains. The results of low-temperature fluorescence emission spectra and chlorophyll fluorescence showed that recombinant phycocyanin has considerable optical activity. The expression of phycocyanin, heme oxidase and ferredoxin oxidoreductase in low-light conditions is particularly evident in the promotion of photosynthesis in *C. reinhardtii*. The growth of transgenic strains was significantly promoted in the early growth phase under low-light conditions. However, the final growth and biomass accumulation of transgenic *C. reinhardti* were inhibited by this expression. In this paper, the possibility of photoenergy transfer between phycocyanin and heterologous host thylakoid membrane was researched, which provided a useful attempt for the construction of a new photosynthetic system using phycobiliprotein from cyanobacteria.

**One-sentence summary:** Phycocyanin from *Arthrospira platensis* FACHB 314 expressed in *Chlamydomonas reinhardtii* can effect the photosynthetic system of *C. reinhardtii*.

---

*Chlamydomonas reinhardtii* is a model organism in the modern research of microalgae. This unicellular green alga offers a simple life cycle and easy isolation of mutants (Harris, 2001). There are a lot of mature tools and techniques for molecular genetic studies with *C. reinhardtii* (Griesbeck et al., 2006). In recent years, reports on the use of microalgae *C. reinhardtii* in studies of biofuels and wastewater treatment are not uncommon (Pittman et al., 2011; Kothari et al., 2013). Some scholars have done some excellent studies on photosynthesis and radiosity of *C. reinhardtii* (Murakami, 1997; Tetali et al., 2007). There are also researchers who refer to *C. reinhardtii* as a most promising cell factory (Johanningmeier and Fischer, 2010). They have expressed lots of pharmaceutical and biotechnological proteins with the *C. reinhardtii* expression system (Franklin and Mayfield, 2004). Moreover, the photosynthetic pigment on the thylakoid membrane in *C. reinhardtii* is similar to that of higher plants. So *C. reinhardtii* is suitable for studying photoenergy transfer of thylakoid membrane through gene transformation and recombination.

Photosynthetic cyanobacteria generally contain chlorophyll a only, and accessory phycobiliprotein pigments which consisting of covalently linked chromophore-protein are the key to enhancing light capture (Lauceri et al., 2018). Phycocyanin is one of the most common phycobiliproteins, which consists of two subunits (β and α). An important step in phycocyanin biosynthesis is the covalent attachment of phycobilin chromophores to specific cysteine (Cys) residues within apo-phycocyanin through a thioether bond (Sun et al., 2019). So phycocyanin has a brilliant color and light absorption properties (Biswas et al., 2010; Zhang et al., 2015). The biosynthesis of phycocyanin is based on heme, and then produced by the reduction and isomerization steps catalyzed by heme oxidase (Ho) and ferredoxin oxidoreductase (PcyA) (Frankenberg et al., 2001). Therefore, these three genes, the apo-phycocyanin gene (*cpcBA*), heme oxidase gene (*ho*) and ferredoxin oxidoreductase (*pcyA*), are crucial in the synthesis of optically active cyanobacteria.

Su et al. successfully expressed allophycocyanin β subunit and allophycocyanin in succession in *C. reinhardtii*. Both of these two foreign recombinant protein which expressed in *C. reinhardtii* transformants accounted for about 2% to 3% of total soluble proteins of the respective strains, respectively (Su et al., 2005; Su et al., 2006). Kenneth et al. have expressed the heterologous *Dunaliella tertiolecta* fatty acyl-ACP thioesterase into *C. reinhardtii*, and the results indicate that *C. reinhardtii* transformants have produced 63% and 94% more neutral lipids than the wild-type. These transformants have an approximately 56% improvement in total lipids without compromising growth (Tan and Lee, 2017). Kwon et al. overexpressed the *FAB2* gene which encodes stearoyl-acyl carrier protein desaturase in *C. reinhardtii* by nuclear transformation, and obtained a mutant strain which has an oleic acid (18:1) content increased by about 2.4-fold compared to the wild-type control plant (Hwangbo et al., 2014). In a word, all of these outstanding work demonstrates that the expression of a fluorescent phycocyanin in *C. reinhardtii* is feasible.

In this study, we attempted to detect the expression of phycocyanin gene in *C. reinhardtii* by nuclear transformation. The phycocyanin gene (*cpcBA*), heme oxidase gene (*ho*) and ferredoxin oxidoreductase gene (*pcyA*) encoding phycocyanin, Ho and PcyA protein was cloned from *Arthrospira platensis* FACHB 314. The transformants were used to study the possibility of photoenergy transfer between phycocyanin and heterologous host thylakoid membrane, which provided a useful attempt for the construction of a new photosynthetic system using phycobiliprotein from cyanobacteria.

## Results

### Generation of transgenic lines of *C. reinhardtii* expressing phycocyanin, Ho and PcyA

To study the expression of phycocyanin genes in *C. reinhardtii* in this research, transgenic lines were created in which genes encoding phycocyanin, Ho and PcyA from *Arthrospira platensis* FACHB 314 were introduced into the nuclear genome. Positive colonies were selected from plates and verified by PCR with primers for *cpcBA*, *ho*, *pcyA* and *aph7*”, which were called strain PC and PCHP containing plasmid pHyg3-*cpcBA* and pHyg3-*cpcBA*-*ho*-*pcyA*, respectively. And the control strain CC-849 is abbreviated as cc849. As shown in the Fig. 1, the *cpcBA*, *ho*, *pcyA* and partial *aph7*” genes with sequence lengths of 1119 bp, 729 bp, 744 bp and 1018 bp were cloned and displayed on an agarose gel electrophoresis map. The templates for the test were from the transgenic lines PC, PCHP and the control cc849. The results showed that no bands of exogenous genes *cpcBA*, *ho*, *pcyA* and *aph7*” were found in cc849. *CpcBA* and *aph7*” genes were cloned clearly from the transformant PC. The bands of *cpcBA*, *ho*, *pcyA* and *aph7*” were cloned clearly from PCHP. The transgenic lines with correct band were further verified by sequencing and then used for the next experiments.

**Fig.1.**
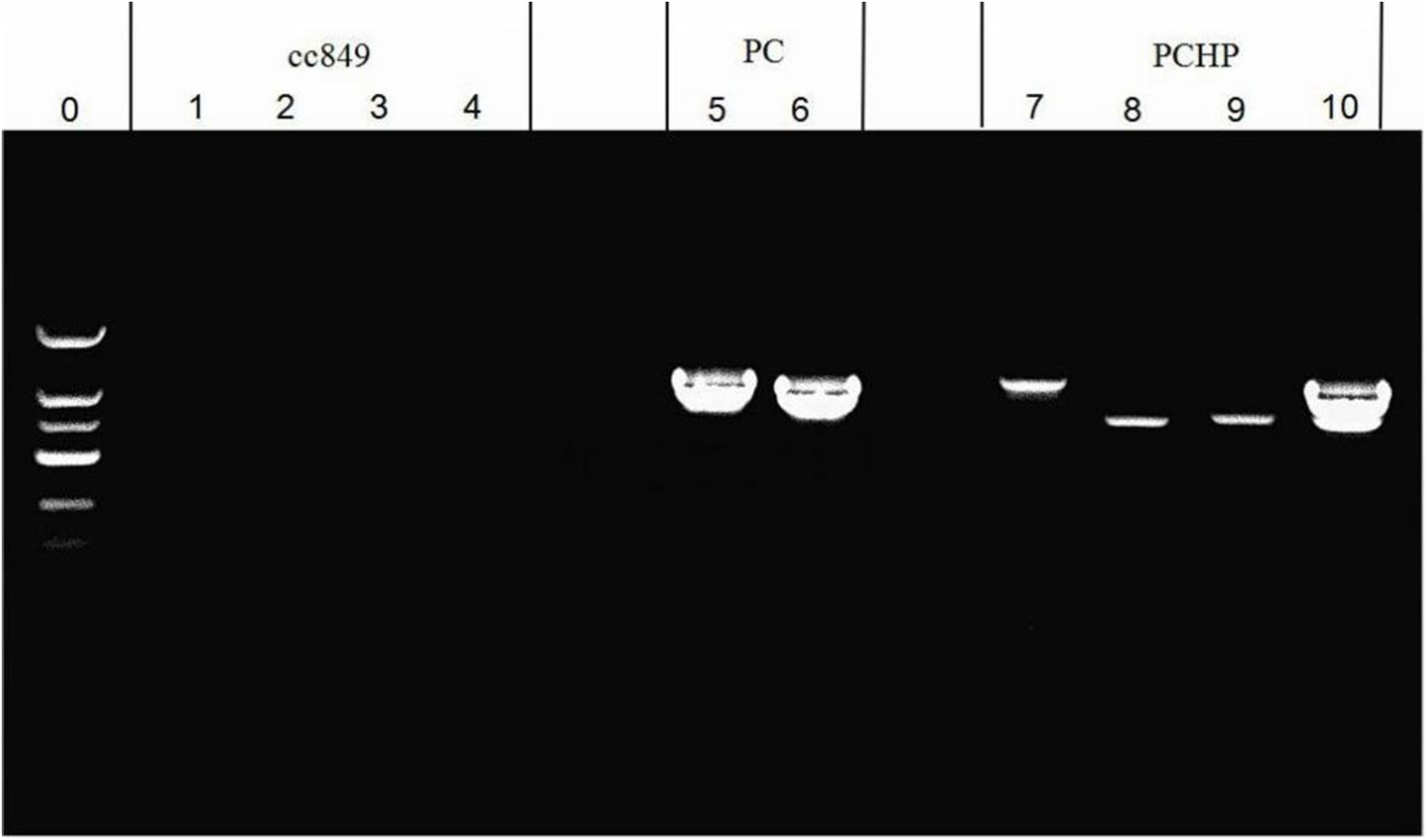
PCR screening of putative transformants to determine their positive.

DNA Marker DL2000 is in the lane 0. Lane 1, 2, 3 and 4 show the PCR result of cc849. No bands of exogenous genes *cpcBA*, *ho*, *pcyA* and *aph7*” were found in cc849. Lane 5 and 6 show the PCR result of the transformant PC with primers of *cpcBA* and *aph7*”. Clear bands indicate the presence of *cpcBA* and *aph7*” genes in the transformant PC. In lane 7, 8, 9 and 10, the bands of *cpcBA*, *ho*, *pcyA* and *aph7*” are cloned clearly from PCHP.

### Analysis of *cpcBA*, *ho*, *pcyA* transgene integration into the nuclear genome

To determine whether the *cpcBA*, *ho* and *pcyA* gene cassettes had integrated in these selected transformants, gDNA were extracted from them and southern blotting was performed. The complete *cpcBA*, *ho* and *pcyA* sequences were amplified by high-fidelity DNA polymerase to be used as the probes.

Genomic DNA from both of transgenic lines PC and PCHP was digested with both of these two group restriction enzymes *Apa*I/*Sal*I and *EcoR*I/*Sac*I. As shown in the Fig. 2, for *cpcBA* gene, when the gDNA was digested with *Apa*I and *Sal*I, there were two hybridization bands for PC and one band for PCHP. When the transformants gDNA digested with *EcoR*I and *Sac*I, it showed clearly one band for both PC and PCHP. This indicates that the exogenous gene *cpcBA* was successfully integrated into genome of the transgenic strain PC and PCHP. And for *ho* and *pcyA* genes, there is one clear band presented in the transformant PCHP. At the same time, there is no blot in lanes of untransformed control strain cc849. This indicates that the exogenous gene *cpcBA* was successfully integrated into genome of the transgenic strain PC as well as *cpcBA*, *ho* and *pcyA* were integrated into the strain PCHP.

**Fig. 2.**
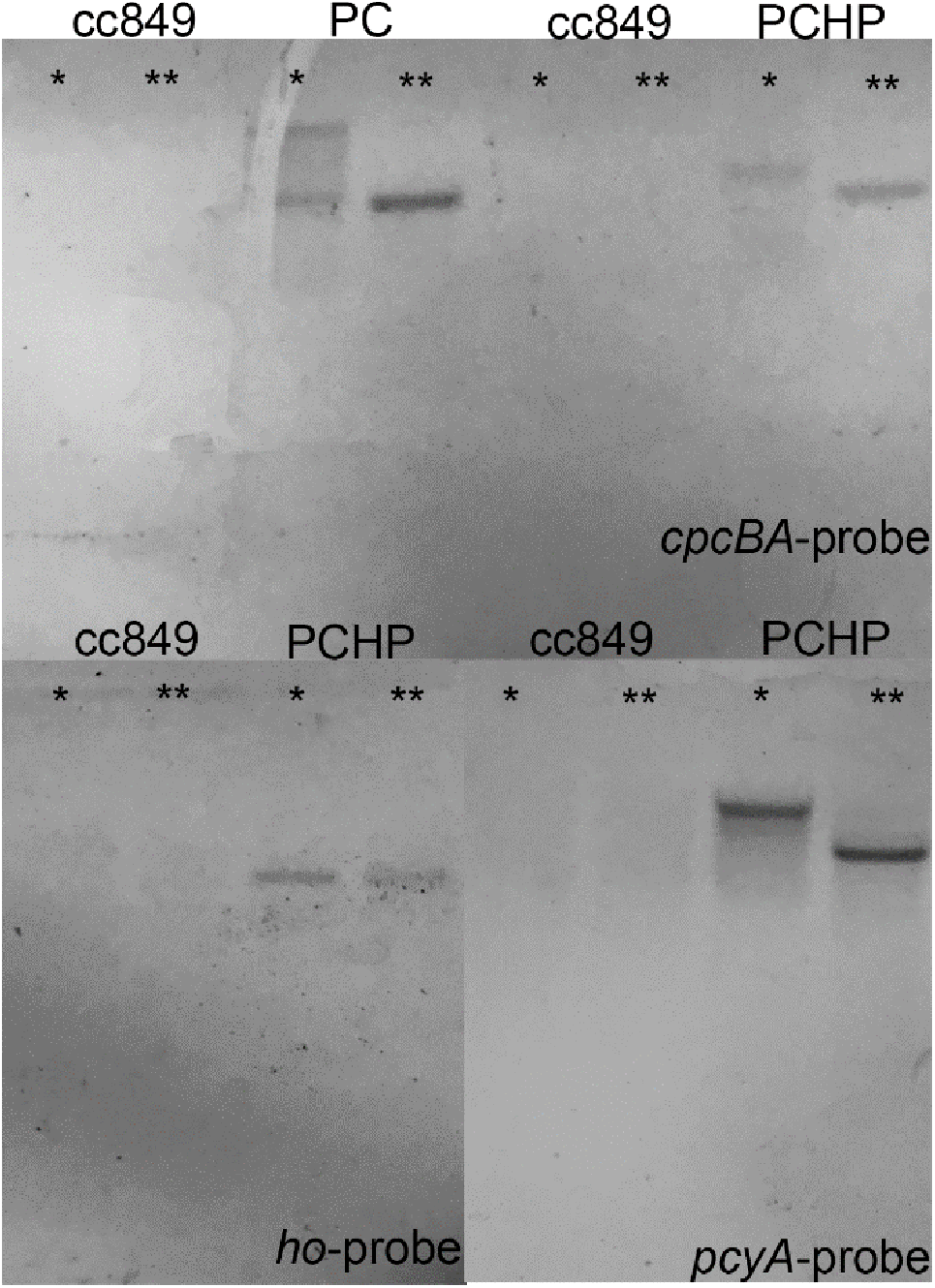
Southern blot for cc849, PC and PCHP. The southern blot analysis was carried out using probes of *cpcBA*, *pcyA* and *ho*, respectively. The samples were gDNA derived from cc849, PC and PCHP. Wherein the symbol ‘*’ represents digestion of the sample with *Apa*I and *Sal*I. And the symbol ‘**’ represents digestion of the sample with *EcoR*I and *Sac*I.

### Fluorescence emission spectra at 77 K

In order to determine the optical activity of recombinant phycocyanin and the effect on the photosystem of *C. reinhardtii*, the tranformed strains of PC, PCHP and the control cc849 cultured under 25 μmol photon m^−2^ s^−1^ (low-light) and 50 μmol photon m^−2^ s^−1^ (high-light) were used to measure low-temperature fluorescence emission spectra at excitation of 435 nm (for Chlorophyll a) and 580 nm (for phycocyanin). As shown in Fig. 3, after excitation at 435 nm, PC and PCHP showed a fluorescence emission peak at 670 nm that was not found in cc849. In low-light experiment, PCHP’s peak at 670 nm is higher than PC (p=2.4111E-25), and its emission peaks representing PS II and PS I (685 nm and 715 nm) (Lapaille et al., 2010) are much higher than PC and the control cc849 (Fig. 3A) (p=2.29674E-13 and 4.08275E-29). At the position of the characteristic peak of phycocyanin (620 nm), PC and PCHP have a small fluorescence emission peak. However, the peak value of PC is higher than that of PCHP (p=1.34327E-50), which may be due to the more energy transfer from phycocyanin to the photosynthetic system in PCHP, resulting the decrease of fluorescence intensity of phycocyanin in PCHP. In the high-light experiment, the peaks of fluorescence of PCHP and PC at 670 nm were higher than that of the control cc849 (p=1.95662E-47 and 1.87227E-67), and the peak of PC is higher than that of PCHP (p=0.008629299), but the difference between these two transgenic lines was not as significant as that in low light conditions (Fig. 3C). The fluorescence emission peaks of PCHP at 685 nm and 715 nm are slightly higher than PC and cc849 (p= 1.4042E-05 and 2.57491E-12), and the differences are still significant. The result means that the recombinant phycocyanin in both PC and PCHP has a positive effect on the photosynthetic system of *C. reinhardtii*, which is more pronounced in low-light conditions.

**Fig. 3.**
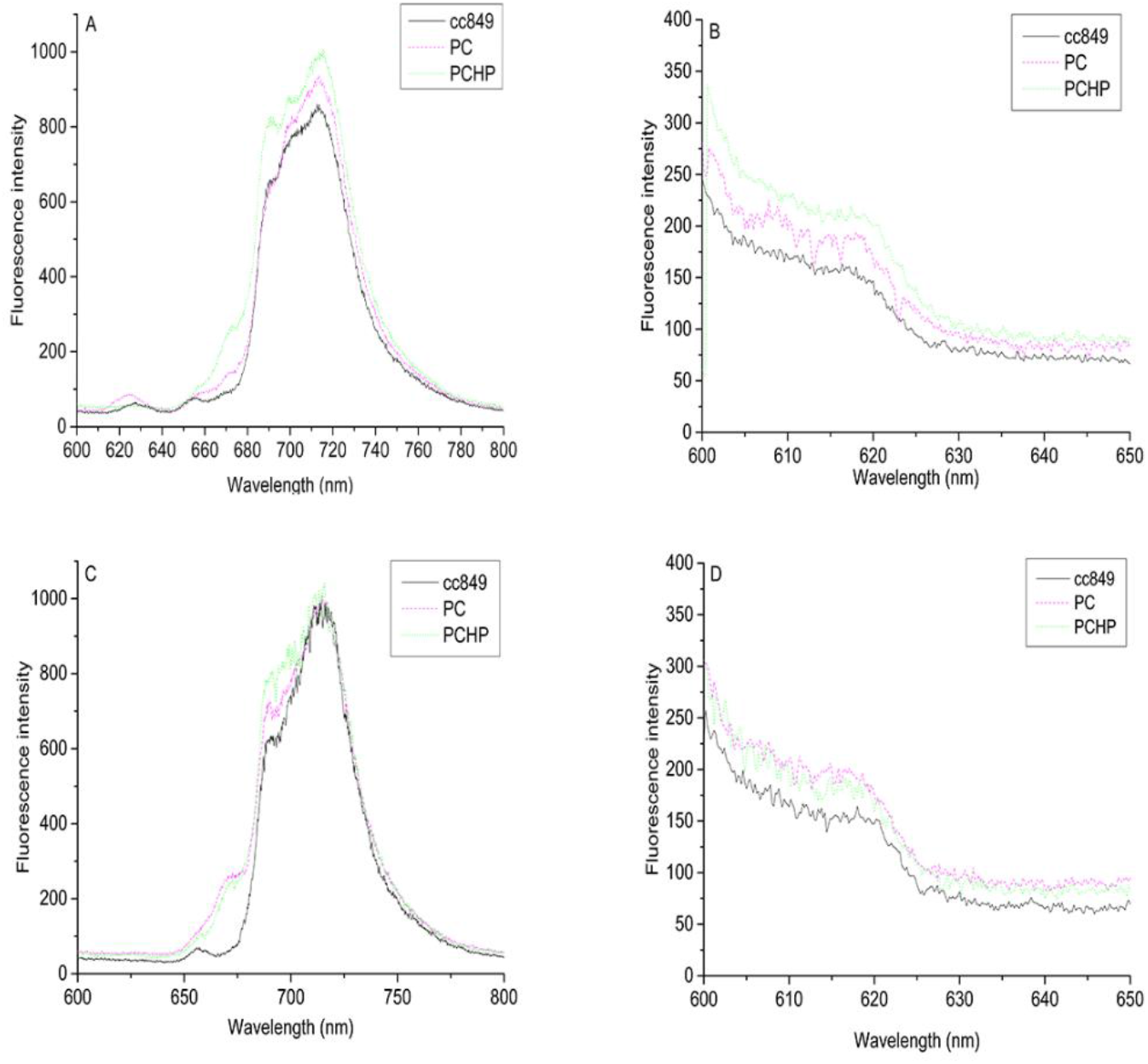
Fluorescence emission spectra of the strains cc849, PC and PCHP. **A.** The emission spectra of these three strains after excitation at 435 nm in the low-light experiment. **B.** The emission spectra of these three strains after excitation at 580 nm in the low-light experiment. **C.** The emission spectra of these three strains after excitation at 435 nm in the high-light experiment. **D.** The emission spectra of these three strains after excitation at 580 nm in the high-light experiment.

When the sample was excited by the characteristic excitation of phycocyanin at 580 nm, in the low-light experiment, the fluorescence emission values of cc849, PC and PCHP at 620 nm were successively increased (Fig. 3B) (p= 1.03818E-06 and 2.18922E-19). While in the high-light experiment, the fluorescence emission values of PCHP and PC at 620 nm are not much different (p= 0.139672), but both of them are higher than the control cc849 (Fig. 3D) (p=2.17031E-08 and 5.98448E-06). This indicates that recombinant phycocyanin in both PC and PCHP has fluorescence activity. And the phycocyanin in PCHP present stronger fluorescence activity under low-light condition.

### Transcription level analysis

RNA of transgenic and control *C. reinhardtii* on days 0, 3, 6, and 9 was extracted for transcription analysis. The transcription level of *cpcBA* was analyzed in PC, PCHP and the untransformed stain cc849. For PCHP, the transcription levels of *ho* and *pcyA* were also analyzed. *RACK* 1 as the housekeeping gene was used to be an internal reference gene for transcriptional level analysis.

The result is shown in the Fig. 4. All transcriptional levels were calculated using fluorescence quantitative values of the corresponding genes in the control as a reference. In the low-light experiment, *cpcBA* in PC and PCHP strains were transcribed successfully, and the transcription level of PCHP was higher than that of PC (p= 0.001651), but both of them showed a downward trend with time (Fig. 4A). Therefore, cells may accumulate more phycocyanin in the early stage, and PCHP may accumulate more phycocyanin than PC. At the same time, *ho* and *pcyA* also transcribed successfully in the transformant PCHP and the transcription levels decreased gradually (or remained roughly unchanged) with the culture time (Fig. 4B, 4C). Because of these, the transgenic strains may express more optical active phycocyanin in the early growth in the low-light condition. On the other hand, in the high-light experiment, *cpcBA* in PC and PCHP strains were transcribed successfully, but the transcription levels of *cpcBA* in PC gradually increased, while in PCHP decreased on the 9^th^ day (Fig. 4D). In addition, the transcription levels of *ho* and *pcyA* in PCHP increased first and then decreased slightly, but in general, the expression level remained at a high level all the time. (Fig. 4E, 4F).

**Fig. 4.**
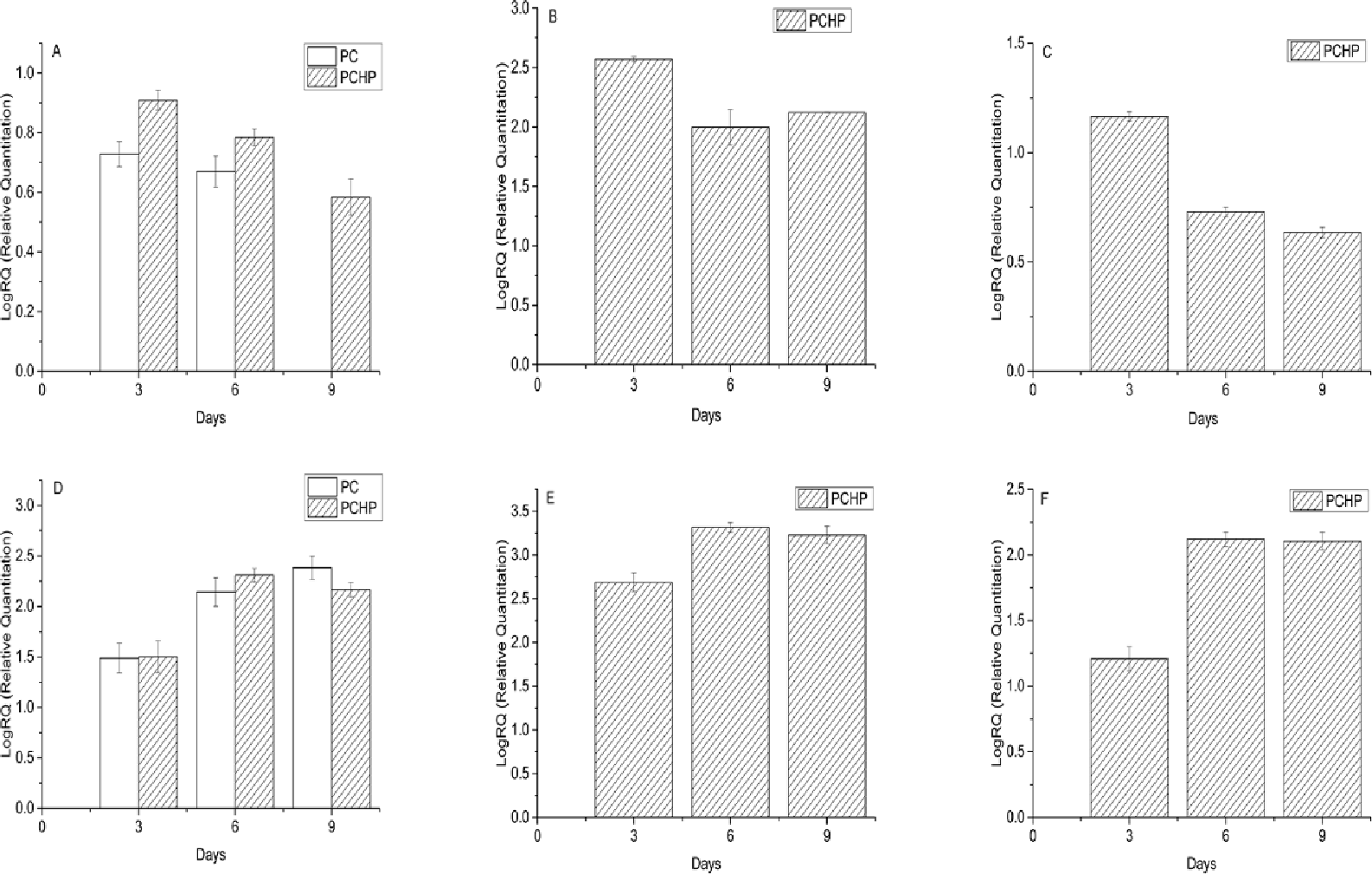
Transcriptional level analysis of transgenic *C. reinhardtii*. **A.** Transcriptional level of *cpcBA* in strains PC and PCHP under low-light conditions. **B.** Transcriptional level of *ho* in strain PCHP under low-light conditions. **C.** Transcriptional level of *pcyA* in strain PCHP under low-light conditions. **D.** Transcriptional level of *cpcBA* in strains PC and PCHP under high-light conditions. **E.** Transcriptional level of *ho* in strain PCHP under high-light conditions. **F.** Transcriptional level of *pcyA* in strain PCHP under high-light conditions.

By comparison, transgenic *C. reinhardtii* expressed more phycocyanin under high-light conditions. It has been reported that the expression of *cpcB* in *Arthrospira platensis* is affected by the light intensity under the action of its upstream regulatory elements (Lu and Zhang, 2005). However, since we have not introduced the relevant regulatory elements into *C. reinhardtii*, this difference in expression may be due to the fact that higher light intensity could provide cells more energy and carbon for life activities, such as protein synthesis. So under low-light conditions, transgenic strains mainly expressed phycocyanin in the early stage of growth. And under high-light conditions, the expression level of phycocyanin was higher and the expression time was longer.

Combined with fluorescence emission analysis, the transcription level of PCHP is higher than PC at low light intensity, which is consistent with its higher fluorescence intensity at low light intensity. Under high-light condition, the transcription level of PCHP and PC are not very different. It may be because of this that the difference in fluorescence peaks between these two is not as significant as under low-light conditions.

### The changes of total chlorophyll content in the transformed *C. reinhardtii*

Intracellular chlorophyll levels are important physiological index of photosynthesis (Zhou et al., 2016). Phycocyanin and chlorophyll are both photosynthetic pigments. In order to study whether the expression of phycocyanin affects the content of chlorophyll, the trend of total chlorophyll content with time in each strain was researched (Fig. 5). The total chlorophyll content in cc849, PC and PCHP was named as TChlcc_849_, TChl_PC_ and TChl_PCHP_, respectively. In the low-light experiment, TChlPCHP was always lower than TChl_cc849_ and TChl_PC_. TChl_PC_ was higher than TChl_cc849_ on the 4^th^ day and then below it. In the high light-experiment, TChlPCHP was lower than TChl_cc849_ and TChl_PC_ in the first 6 days. TChlPC was only higher than TChl_cc849_ on the 4^th^ day. The decrease of chlorophyll content in PCHP was presumed to be due to the expression of Ho and PcyA. Phycocyanobilin makes phycocyanin optically active. Chlorophyll and phycocyanobilin are derived from the common precursor protoporphyrin IX (Willows, 2019). Under the catalysis of Ho and PcyA in the transformant PCHP, the protoporphyrin IX is competed to synthesize phycocyanobilin, resulting in a decrease in chlorophyll content. In the transformant PC, chlorophyll content is almost the same as the control cc849, the difference is not significant, which indicates that just the expression of phycocyanin has no obvious effect on the content of chlorophyll.

**Fig. 5.**
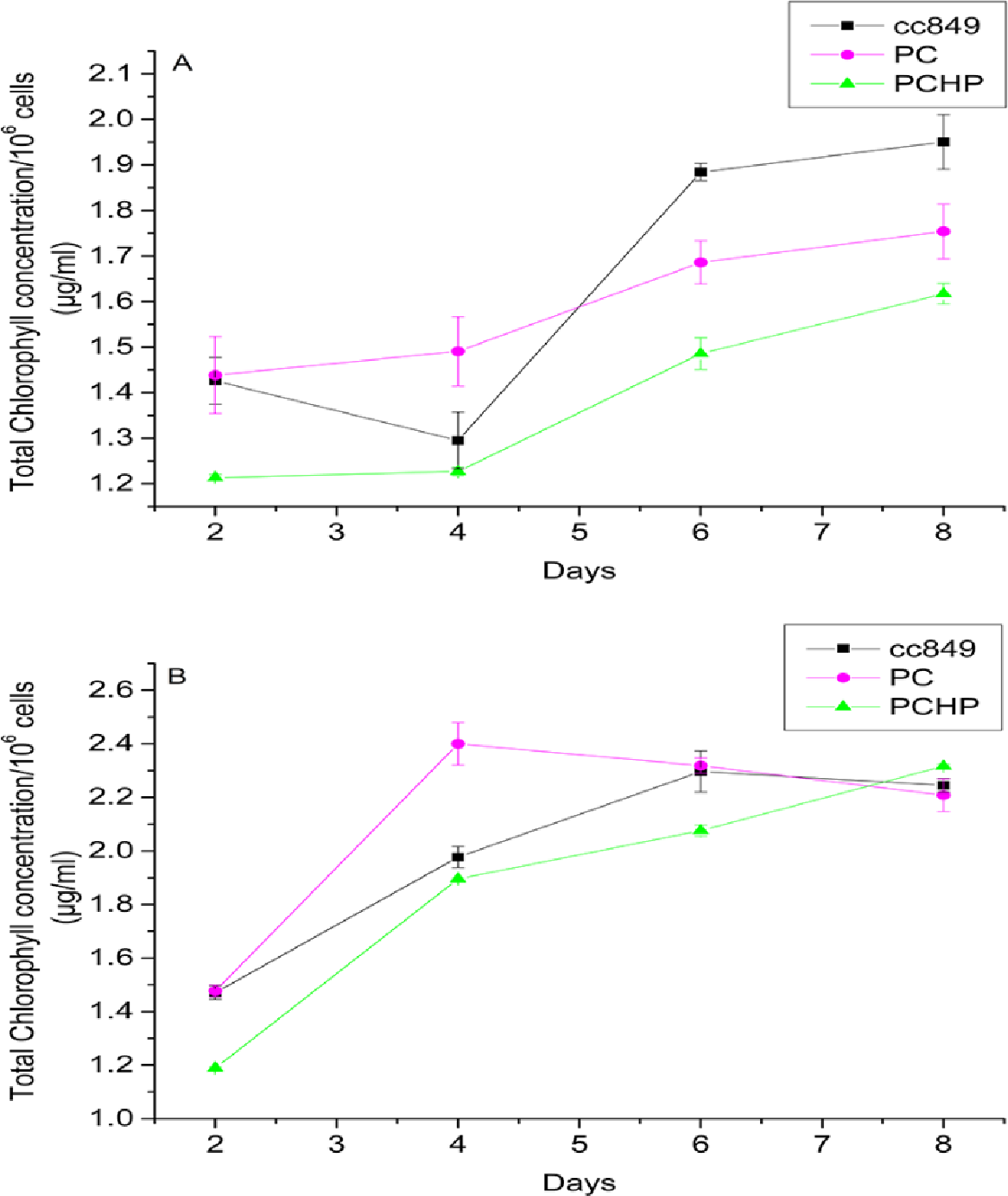
Total chlorophyll content of these three strains (the number of cells was normalized to 10^6^ cells). **A.** Total chlorophyll content in cc849, PC and PCHP in low-light conditions. **B.** Total chlorophyll content in cc849, PC and PCHP in high-light conditions

### Effect of recombinant phycocyanin on PSI and PSII of *C. reinhardtii*

For samples from the transgenic and control *C. reinhardtii* of the 2^nd^, 4^th^, 6^th^ and 8^th^ day, the apparent electron transport rate of PS I and PS II (ETR_PS I_ and ETR_PS II_) were measured and recorded simultaneously. In the low-light experiment, on the 2^nd^ day, the ETR_PS I_ of PC and PCHP was similar in value (p= 0.538921) and higher than that of the control cc849 (p=0.015745 and 0.049245) (Fig. 6A). On the 4^th^ day, the ETR_PS I_ of the PC and the cc849 were almost the same (p= 0.65474), while that of the PCHP was lower than them (p= 0.028189 and 0.046256) (Fig. 6B). On the 6^th^ day, ETR_PS I_ of PCHP was almost the same with cc849 (p= 0.444449), while PC remained higher than PCHP and cc849 (p=0.014856 and 0.047848) (Fig. 6C). On the 8^th^ day, the ETR_PS I_ of PC, cc849 and PCHP were successively decremented (p= 0.041302 and 0.001572) (Fig. 6D). Generally speaking the ETR_PS I_ of PC always remained at a relatively high level, while the ETR_PS I_ of PCHP was higher than cc849 only in the first two days and lower than cc849 in the latter.

**Fig. 6.**
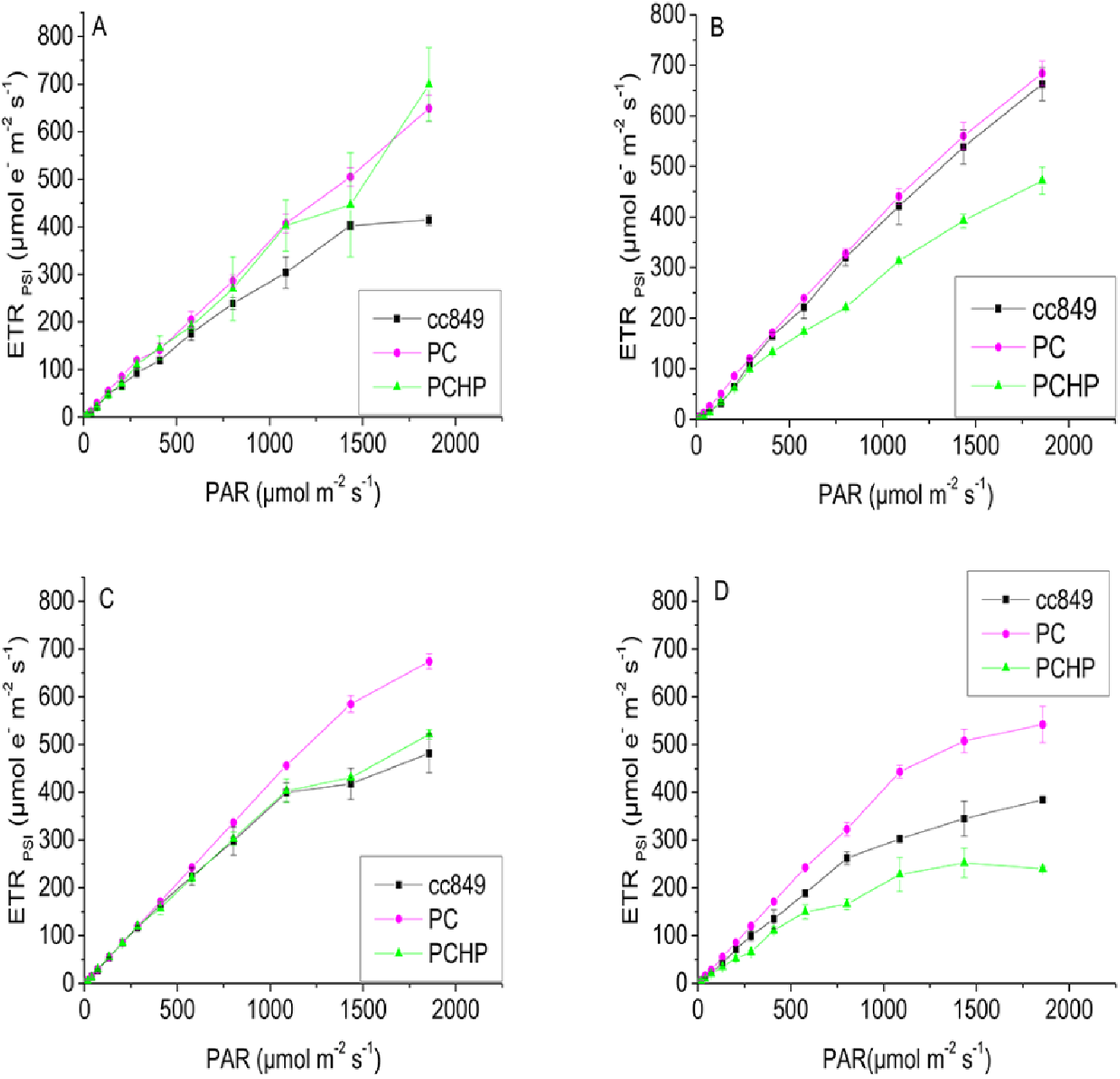
Apparent electron transport rate of PS I in low-light experiment. **A**, **B**, **C**, and **D** The apparent electron transport rate in PSI (ETR_PS I_) in cc849, PC and PCHP on the 2^nd^, 4^th^, 6^th^ and 8^th^ day.

For ETR_PSII_, PCHP and PC was higher than cc849 on the 2^nd^ day (p= 0.012463 and 0.026164) (Fig. 7A). On the 4^th^ day, the ETR_PS II_ of cc849 was higher than that of PCHP and PC (p= 0.0495761 and 0.014351) (Fig. 7B), but the difference was not significant. And then on the 6^th^ day, ETR_PS II_ of PC were higher than that of cc849 and PCHP (p= 0.041116 and 0.035448) (Fig. 7C). On the 8^th^ day, ETR_PS II_ of these three strains were about the same size (p= 0.857638, 0.970591 and 0.653982) (Fig. 7D). On the whole, the ETR_PS II_ was also affected by the recombinant phycocyanin, especially in the first two days of cultivation.

Overall, the expression of recombinant phycocyanin has an impact on the electron transport rate of photosynthetic system of transformant *C. reinhardtii*, especially on photosystem I. The strain PCHP was significantly positively affected only in the first two days, while PC still received the great influence in the later stages of growth.

**Fig. 7.**
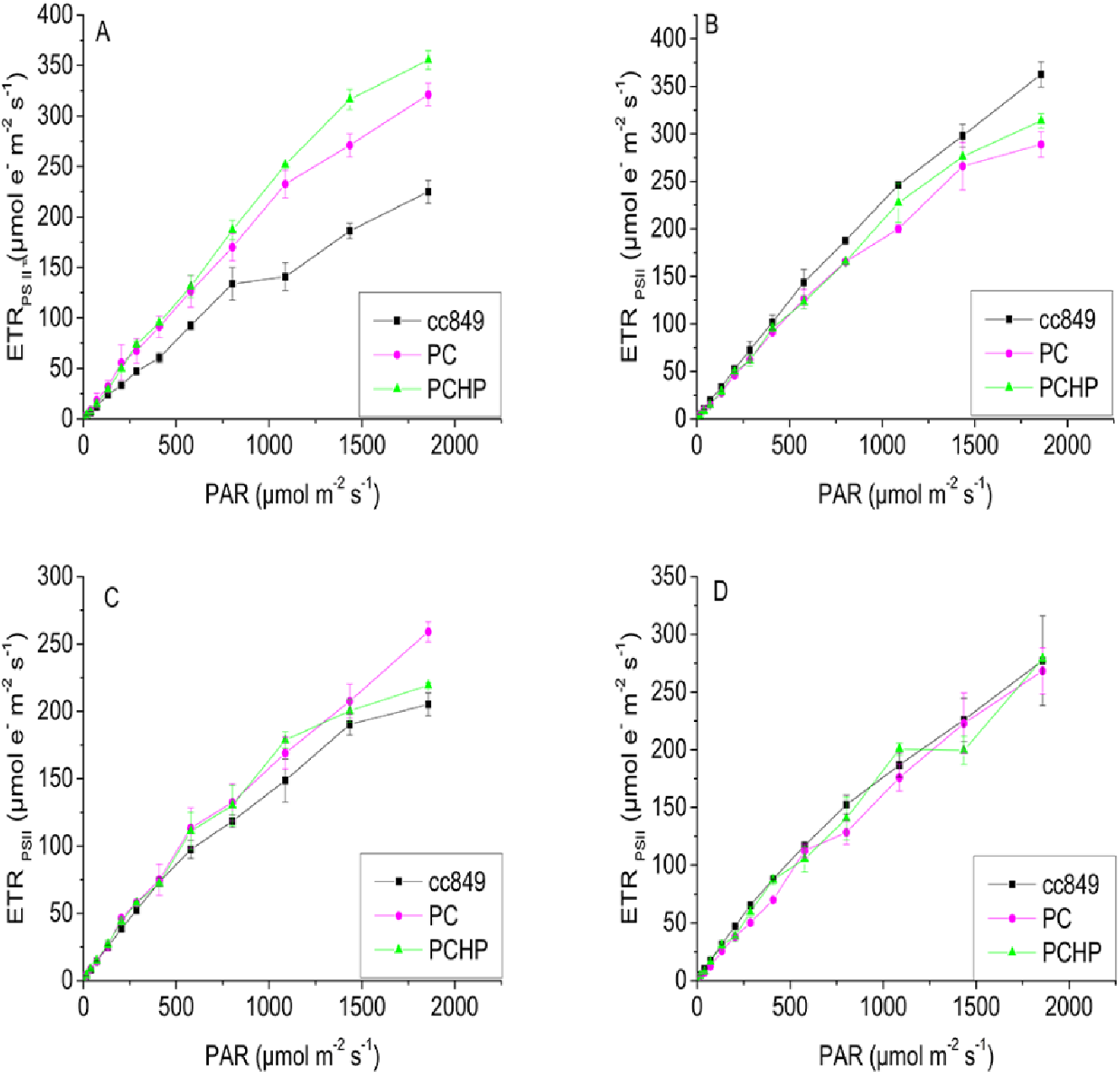
Apparent electron transport rate of PS II in low-light experiment. **A**, **B**, **C**, and **D** The apparent electron transport rate in PSII (ETR_PS II_) in cc849,PC and PCHP on the 2^nd^, 4^th^, 6^th^ and 8^th^ day.

At the same time, in the high-light experiment, ETR_PS I_ of PC and PCHP are higher than cc849 on the 2^nd^ day (p= 0.042833 and 0.033793) (Fig. 8A). But on the 4^th^ day, ETR_PS I_ of cc849 was higher than that of PC and PCHP (p= 0.005064 and 0.003692) (Fig. 8B). And then on the 6^th^ day, PCHP was the highest in the curve of ETR_PS I_ (p= 0.028004 and 0.015244) (Fig. 8C). ETR_PS I_ of PC was higher than cc849 and PCHP on the 8^th^ day (p= 0.01271 and 0.049476) (Fig. 8D). In the high-light experiment, ETR_PS I_ was also affected by the recombinant phycocyanin, especially in the transformant PC.

**Fig. 8.**
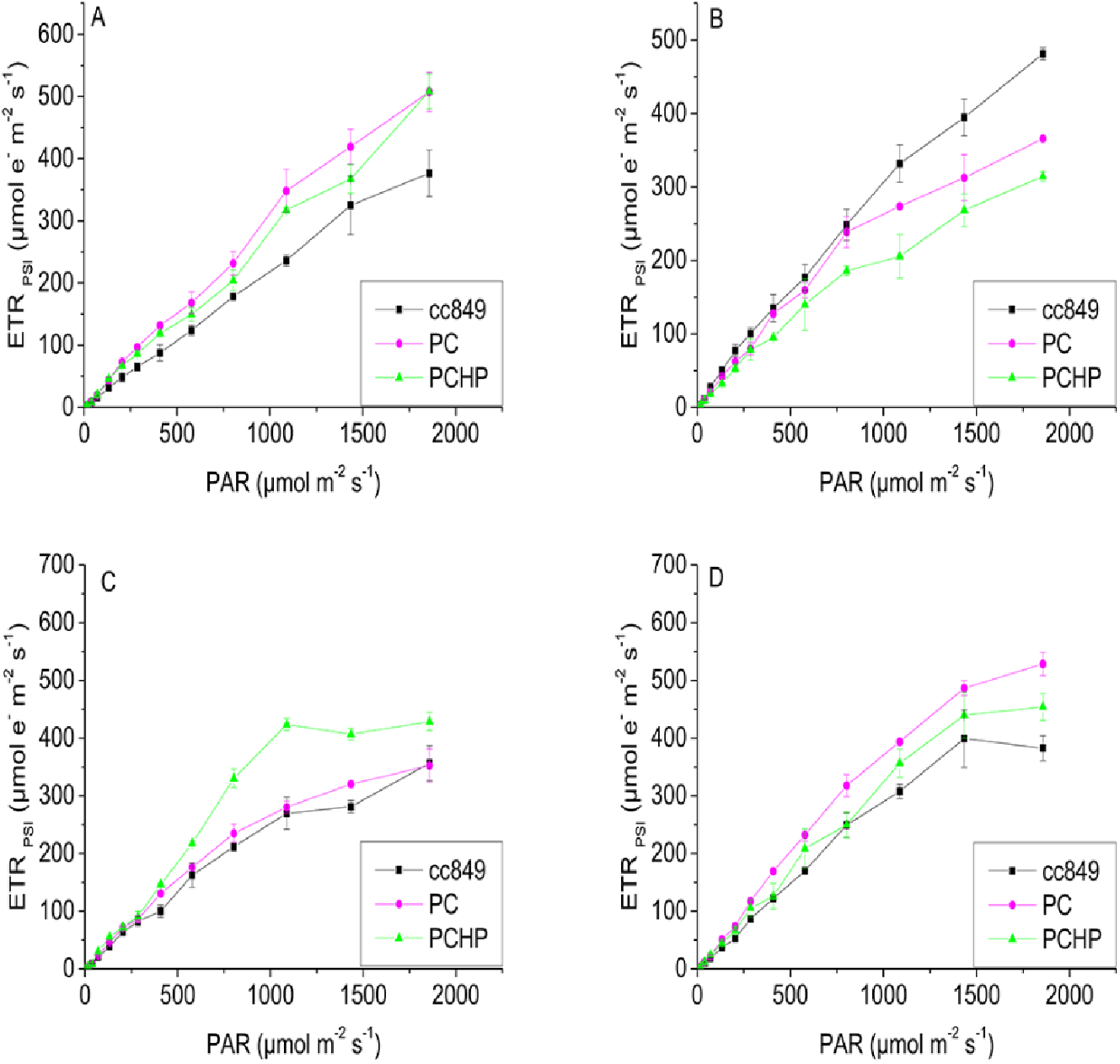
Apparent electron transport rate of PS I in high-light experiment. **A**, **B**, **C**, and **D** The apparent electron transport rate in PSI (ETR_PSI_) in cc849,PC and PCHP on the 2^nd^, 4^th^, 6^th^ and 8^th^ day.

While for ETR_PS II_, at high light intensity, phycocyanin showed little effect. On the 2^nd^ day, these three strains had almost the same ETR_PS II_ (p= 0.768031, 0.79641 and 0.886925) (Fig. 9A). On the 4^th^ day, the ETR_PS II_ of cc849 is even higher than PC and PCHP (p= 0.048283 and 0.0340475) (Fig. 9B). From the 6^th^ to 8^th^ day, ETR_PS II_ of the three strains are similar, and the differences were not significant (Fig. 9C and 9D). In general, at high light intensity, effect of recombinant phycocyanin on the photosynthesis was not obviously.

**Fig. 9.**
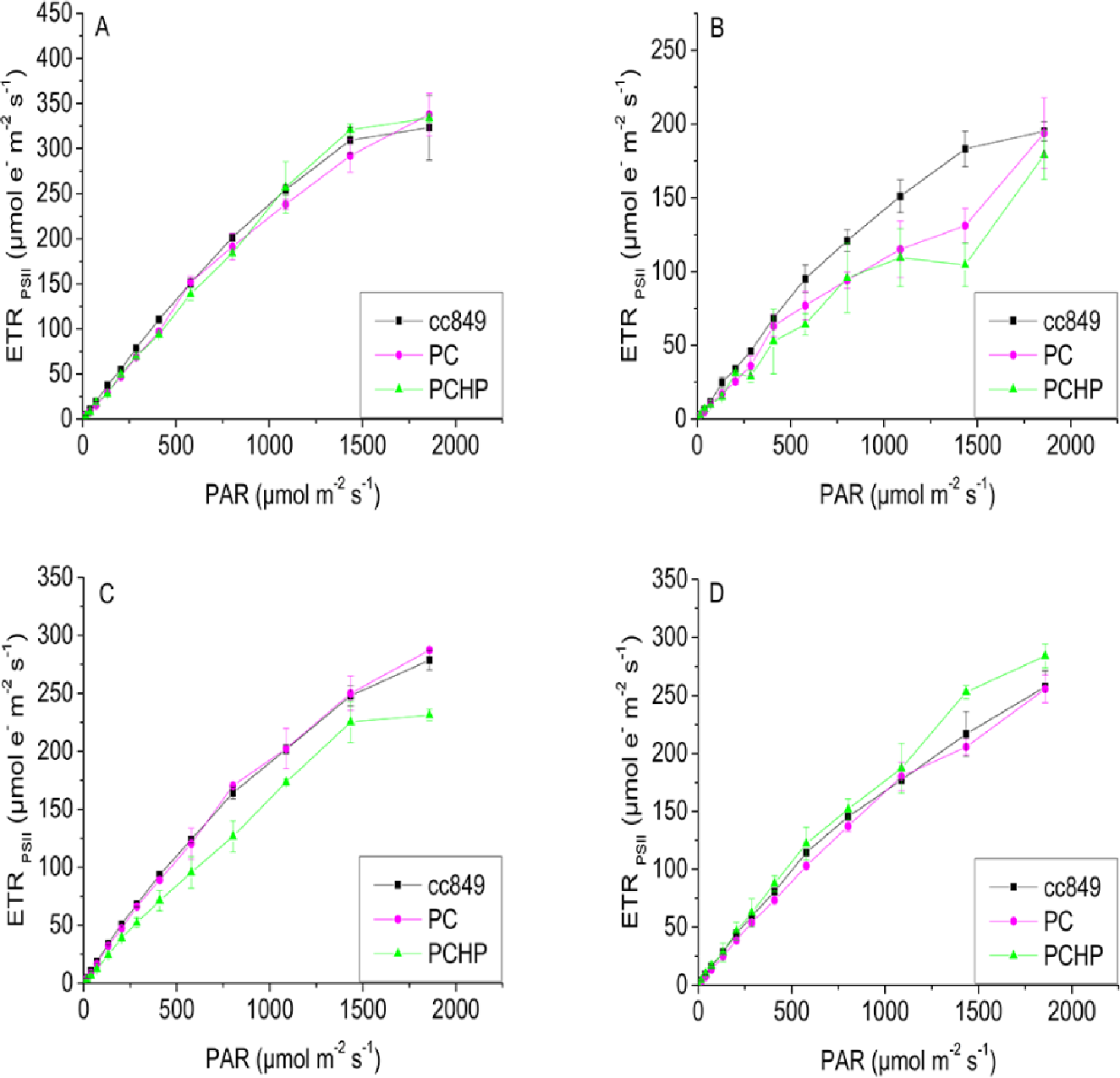
Apparent electron transport rate of PS II in high-light experiment. **A**, **B**, **C**, and **D** The apparent electron transport rate in PSII (ETR_PS II_) in cc849,PC and PCHP on the 2^nd^, 4^th^, 6^th^ and 8^th^ day.

### Effect of expression of phycocyanin, Ho and PcyA on growth and biomass accumulation during mixotrophic growth

To investigate whether transgenic strains can grow faster and/or accumulate more biomass than the control strain cc849, the biomass of *C. reinhardtii* was recorded by measuring OD_750nm_ daily in a 10-day experiment and made a curve. As shown in Fig. 10, in the first 3 days, the cell density of PCHP in the low-light experiment was initially greater than that of the PC and cc849 (p= 0.000283 and 0.000416). However, PCHP entered the stationary phase on the 3^rd^ day, while the latter two strains had longer exponential phase till the 4^th^ day. It is particularly interesting, compared to PCHP, which maintained a stable cell density, the density of PC and cc849 dropped sharply on the 7^th^ day after maintaining a short-term stability for 3 days, followed by a slight increase on the 8^th^ day, and remained stable until the 10^th^ day. This may be due to the exhaustion of the carbon source in the medium, which results in the cells being able to obtain carbon only through photosynthesis. And photosynthesis in low-light conditions could not provide enough light energy for cells to sustain vigorous growth and limit cell growth. After the 4^th^ day, the cell density of PC and cc849 was always greater than that of PCHP (p= 3.91159E-06 and 0.000512). While the density of PC is slightly larger than cc849 (p= 3.91159E-06), which may be due to the more energy captured by the recombinant phycocyanin in strain PC. In the final stationary phase, the densities of PC and PCHP were 107% and 91.2% of cc849, respectively. Dry weight measurements also showed that biomass of PC was slightly higher than that of cc849 without a significant difference (p= 0.175446), but both of them were higher than PCHP (p= 0.01207 and 0.019682). The dry weight of PC and PCHP was 1.23 and 0.47 times that of cc849, respectively. The growth advantage of PCHP in the first three days may indicate that the expression of phycocyanin has a positive effect on the growth of *C. reinhardtii*, while the poor growth in the end may be due to the energetic load of *C. reinhardtii* from the excessive expression of foreign proteins (Dejtisakdi and Miller, 2016).

**Fig. 10.**
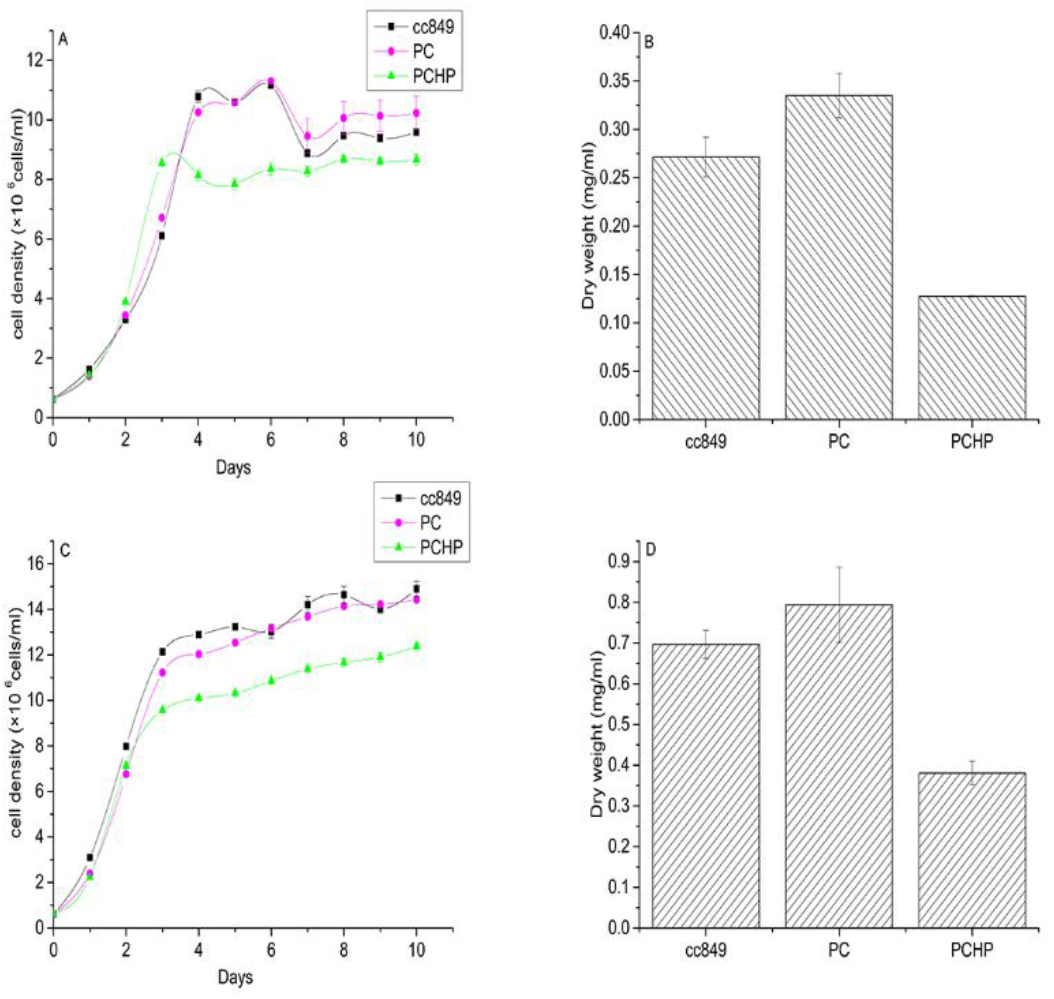
growth and biomass accumulation during mixotrophic growth. **A.** Growth curve of three *C. reinhardtii* strains under low-light conditions **B.** Comparison of dry weight of three strains of *C. reinhardti*i under low-light conditions **C.** Growth curve of three *C. reinhardtii* strains under high-light conditions **D.** Comparison of dry weight of three strains of *C. reinhardtii* under high-light conditions

On the other hand, in the high-light experiment, the three lines grew at a similar rate at the beginning of the exponential phase. However, after entering the stationary phase at the 3^rd^ day, the cell densities of PC and PCHP were 3.3% and 19.1% lower than cc849, respectively. The dry weight of PC is not much different from that of cc849 (p= 0.426844), while PCHP is 45.3% lower than the control (p= 0.01951). This may indicate that sufficient light allows the control strain to reach the growth limit, and in this situation, the effect of phycocyanin does not fully compensate for the energetic load imposed by the expression of the foreign protein in *C. reinhardtii*.

## Discussion

In this study, two transformation vectors of *C. reinhardtii* were constructed and used to determine the effects of phycocyanin on photosynthesis and growth of *C. reinhardtii*. Reports of phycocyanin expression in *E. coli* are not uncommon. For example, Tooley et al. expressed a fluorescent cyanobacterial C-phycocyanin holo-alpha subunit in *E. coli*(Tooley et al., 2001). However, there are not many reports of phycocyanin expression in eukaryotic cells. Our research provided new ideas for improving the growth rate of microalgae and studying the photosynthetic energy transfer in other photosynthetic organisms.

In this research, *cpcBA*, *ho* and *pcyA* from *Arthrospira platensis* were transformed into *C. reinhardtii*. Each of our promoters and terminators in the transformation vector can serve as a homologous integration site and therefore has a strong ability to integrate. Through southern blotting, the single band of *cpcBA*, *ho* and *pcyA* in PCHP demonstrated that they were successfully integrated into the nuclear genome of *C. reinhardtii* by a single copy. While *cpcBA* in PC may be integrated by double copies form. The transcription levels of *cpcBA*, *ho* and *pcyA* in strain PCHP were higher than that of PC in low-light conditions, especially in the early stage. This partly explains that the growth rate of PCHP in the exponential phase in the low-light experimental growth curve is greater than that of PC and cc849.

As shown by the results of the 77K fluorescence spectrum, both PC and PCHP possessed a unique fluorescent peak at 670 nm when compared to the control cc849 as excited at 435 nm. And their peaks representing PSII (685nm) and PSI (715nm) were significantly improved compared to the control. This suggests that energy transfer from phycocyanin to the photosystem may occur in transgenic strains. Wlodarczyk et al. reported the excitation energy transfer of PSII–LHCII/PSI–LHCI and PSI/PSII in intact cells of *C. reinhardtii* in the absence of PSII core or PSI core (Wlodarczyk et al., 2016). It can be seen that the light energy transfer from the light-harvesting antenna to the non-corresponding photosynthetic system core is possible. So this indicates that the energy transfer from phycocyanin to photosynthetic system in *C. reinhardtii* is credible. The peak of PCHP was higher than that in PC in low-light experiments, and these two were not very different in high-light experiments. It suggests that the promotion of phycocyanin to photosynthesis of *C. reinhardtii* is more pronounced in low-light conditions. This was also confirmed by fluorescence at 620 nm when excited at 580 nm in low-light and high-light experiments. And this is consistent with the experimental results of PCHP in the exponential phase at high growth rate compared with PC and cc849, and the growth rate of the three is similar under high-light conditions. In addition, the characteristic peak of phycocyanin was also detected in PC strains without phycocyanobilin synthesis, suggesting that *C. reinhardtii* may have a chromophore similar to phycocyanobilin, which can bind to phycocyanin and give it optical activity. This may be present in view of the fact that some scholars have isolated red pigments from mutant strains of *C. reinhardtii*, which have a structure similar to that of bile pigments (Doi et al., 1997).

Analysis of the low temperature fluorescence spectra revealed that the emission spectra of all strains were closer to state 2, which may be due to the decrease in PQ caused by dark adaptation before measurement (Cardol et al., 2003). Although the state transition of *C. reinhardtii* has the function of regulating the relative speed of photosynthetic linear and cyclic electron flows (Wollman, 2001). Since the control cc849 has undergone dark adaptation for the same time and was also converted to state 2, the elevation of the ETR of the transgenic strain relative to the control may be completely caused by the expression of the foreign phycocyanin. ATP and NAD(P)H in linear electron transport are consumed in the Calvin cycle, and ATP produced by cyclic photophosphorylation can be used in other metabolic processes, so it has a positive effect on cell growth (Cardol et al., 2003). Interestingly, phycocyanin in cyanobacteria and red algae primarily transmits light energy to the PSII (Glazer, 1989; Green et al., 2003). However, our work indicates that phycocyanin had an effect on both of PS II and PS I of *C. reinhardtii*, especially in PS I. This may be because that the separation between the PSI and PSII-rich areas in *C. reinhardtii* is less pronounced here (Wlodarczyk et al., 2016).

According to the analysis of the cell growth curve, the growth rate of PCHP in the exponential phase was higher than that of PC and cc849 under low-light conditions. There were no significant differences between the three under high-light conditions. This also confirmed that the promotion of phycocyanin on *C. reinhardtii* under low-light conditions is more obvious than that in high-light conditions. As for the insignificant difference in growth rate and dry weight between PC and cc849, it may be related to the lack of modification of heme oxidase and ferredoxin oxidoreductase in the phycocyanin with the low activity in PC (Sun et al., 2019). It can be seen that the phycocyanin with fluorescence activity has a considerable positive effect on photosynthesis of *C. reinhardtii*. However, the final growth rate and dry weight accumulation of PCHP were much lower than PC and cc849. This may be due to the fact that the integration of foreign genes impaired cell growth in some way (Dejtisakdi and Miller, 2016). Besides this, this growth disadvantage may be due to the lower total chlorophyll content in PCHP. The expression of HO and PcyA catalyzed the synthesis of phycocyanin from protoporphyrin IX, which resulted in the decrease of protoporphyrin IX and then the decrease of chlorophyll content (Willows, 2019). The decrease of chlorophyll content may also lead to the decrease of growth rate of strain PCHP. Another possibility is that the in vivo expression of a variety of exogenous proteins brings a heavy energetic load to *C. reinhardtii*. Dejtisakdi et.al overexpressed the fructose 1,6-bisphosphatase in *C. reinhardtii* and found a detrimental effect on cells growth from energetic load (Dejtisakdi and Miller, 2016). The expression of phycocyanin in this study may be subject to the same inhibition. In addition, the absence of linker proteins prevents exogenous phycocyanin from anchoring to the thylakoid membrane of *C. reinhardtii* (Tang et al., 2012; Zhang et al., 2017). On the other hand, phycocyanin in algae plays a role on the thylakoid membrane (Sidler, 1994; Liu et al., 2005). But it was integrated into the nuclear genome of *C. reinhardtii* in this work, which may be detrimental to its photoenergy transmission to the thylakoid membrane. Recombinant phycocyanin may be difficult to enter the chloroplast of *C. reinhardtii*. The level of exchange between the two compartments remains an important question (Kumar et al., 2012). It has been reported that light intensity will regulate the expression of metalloproteinase FtsH in the chloroplast (Seo et al., 2000; Wang et al., 2017). So there is also a possible reason that the activity of intracellular proteases may affect the half-life of phycocyanin. In a word, the energy transfer of phycocyanin and photosystem is not efficient enough to compensate for the above energetic load. And for strain PC, the phycocyanin expressed by it is not optically active enough in the absence of phycocyanobilin synthesized by heme oxidase and ferredoxin oxidoreductase. Therefore, the promotion of photosynthesis is far less obvious than PCHP. And these improvements may be completely offset by the energetic load. This is evidenced by its unsuppressed growth compared to the control.

## Conclusion

Phycocyanin, ferredoxin oxidoreductase and heme oxidase from *Arthrospira platensis* FACHB 314 were successfully expressed in *C. reinhardtii*. Our work demonstrated that fluorescently active phycocyanin promotes photosynthesis of *C. reinhardtii*, and this positive effect is more pronounced in low-light conditions. Although the energetic load caused by the expression of foreign proteins and other reasons made the final cell density and dry weight of the transgenic strain with three foreign genes lower than the control strain, this transgenic strain had a greater growth rate in the exponential phase under low-light conditions. And the photosynthesis and growth of PC strain that lack the phycocyanobilin have also been promoted under low-light conditions, compared with the control. This provided an idea for us to increase the growth rate of *C. reinhardtii*, especially under conditions of insufficient light. In addition, our work has laid the foundation for studying the role of cyanobacterial photosynthetic elements in other green-lineage photo-synthetic organisms.

## Materials and methods

### *C. reinhardtii* strain and growth conditions

*C. reinhardtii* cell wall deletion mutant strain CC-849 was obtained from the Chlamydomonas Resource Center, University of Minnesota. The cells were cultured mixotrophically in Tris-acetate-phosphate (TAP) medium, PH6.8, at 23 °C (Chapter 8 - Chlamydomonas in the Laboratory, 2009). Cells cultures were illuminated at 50 (high-light) and 25 (low-light) μmol photon m^−2^ s^−1^ with a 12:12 h dark/light cycle and shaking at 100 rpm daily.

### Plasmid construction

The *C. reinhardtii* transformation vector pHyg3-*cpcBA* and pHyg3*-cpcBA-ho-pcyA* were constructed based on the vector pHyg3 from the Chlamydomonas Resource Center, University of Minnesota. The pHyg3 plasmid was first digested with *Kpn*I and *Nde*I and ligated the cassette consisting of *C. reinhardtii β-2 tubulin* promoter, the phycocyanin gene (*cpcBA*) of *Arthrospira platensis* FACHB 314 and the *C. reinhardtii rbcS*2 gene 3’ untranslated region into it. The restriction sites *AscI*, *Bgl*II and *Mlu*I were introduced to the terminal of the *rbcS*2 3’ untranslated region of the cassette at the same time. Then the *C. reinhardtii* transformation vector pHyg3-*cpcBA* was built successfully. The pHyg3-*cpcBA-ho-pcyA* was constructed based on pHyg3-*cpcBA*. The pHyg3-*cpcBA* plasmid was digested with *Asc*I, *Bgl*II and *Mlu*I, and then the *ho*-cassette and the *pcyA*-cassette were introduced into it respectively, which are composed of *ho*/*pcyA* from *Arthrospira platensis* FACHB 314 flanked by *C. reinhardtii β-2 tubulin* promoter and *C. reinhardtii rbcS2* gene 3’ untranslated region. Then we got the *C. reinhardtii* transformation vector pHyg3-*cpcBA*-*ho*-*pcyA*. The *aph7*” cassette of the original pHyg3 conferred pHyg3-*cpcBA* and pHyg3-*cpcBA*-*ho*-*pcyA* a resistance against hygromycin B (Berthold et al., 2002). Primers used in this study are listed in Table 1.

**Table 1.**
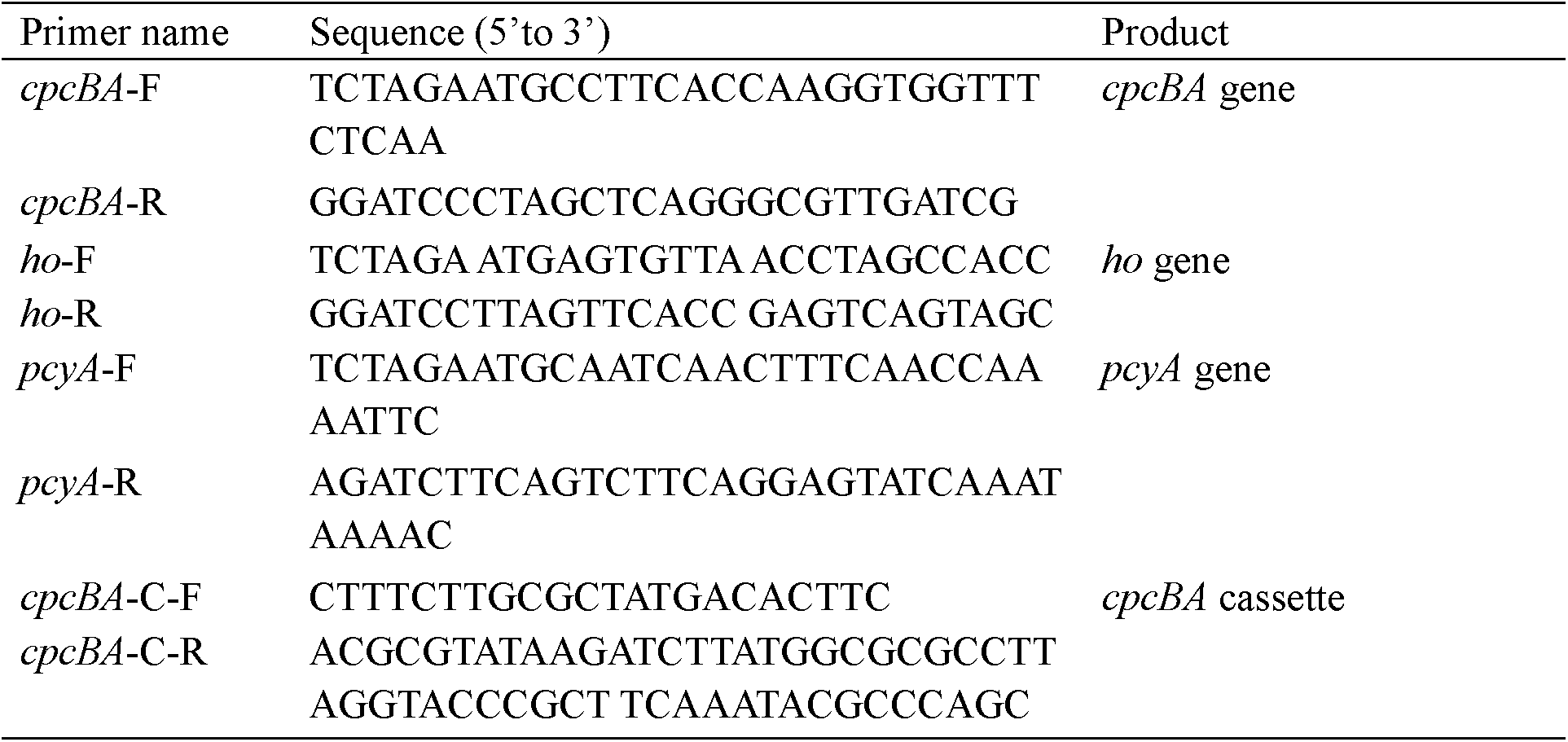
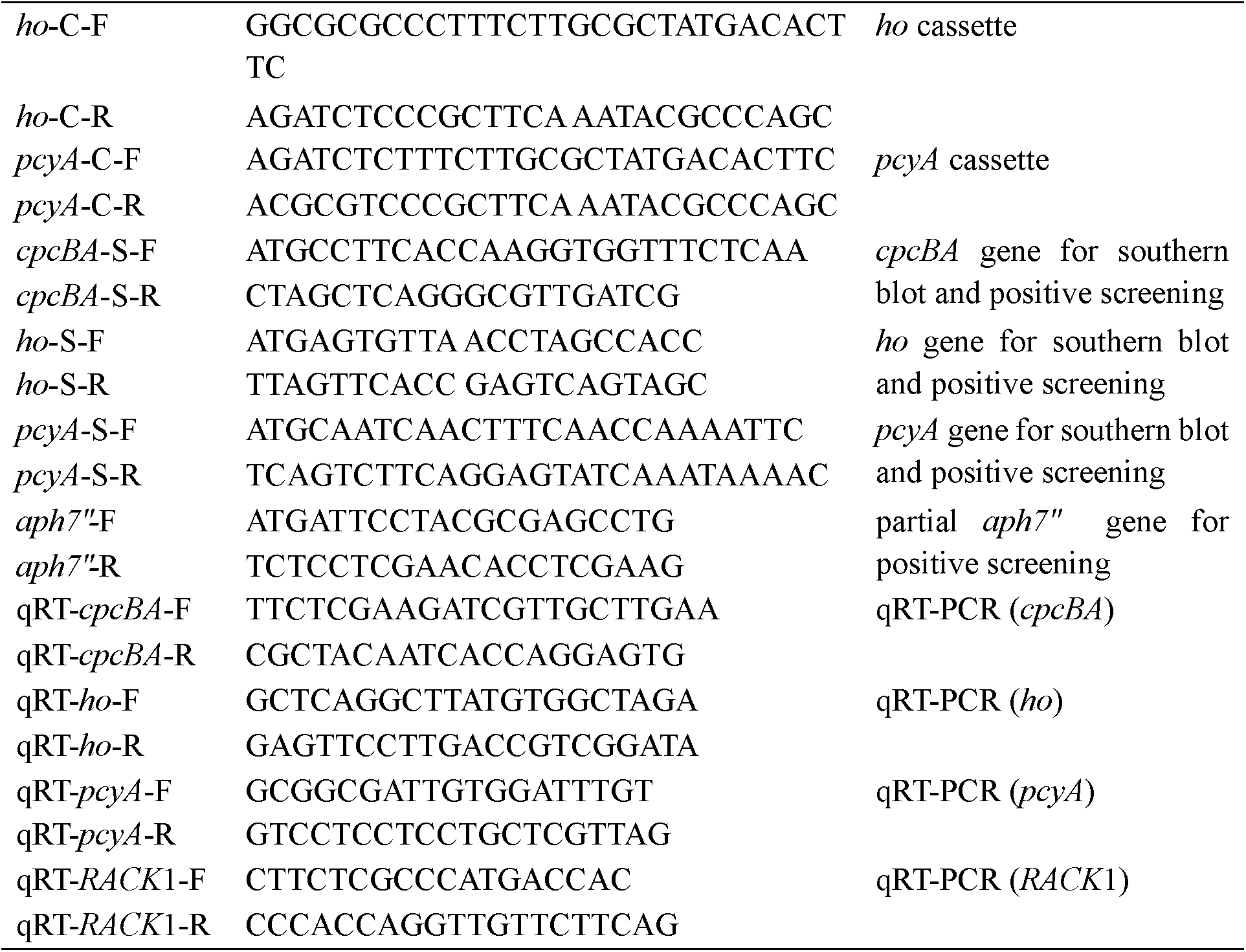
List of primers synthesized in this study

### Transformation of *C. reinhardtii*

The cells of *C. reinhardtii* were prepared for nuclear transformation as follows. The transformation procedure was based on that described by Kindle (Kindle, 1990). Cells were cultured to a cell density of 3–5 × 10^6^ cells/mL. Then cells were harvested by centrifugation at 3000 ×g for 10 min at 20 °C. The pellet was resuspended with TAP medium to a cell concentration of 10^8^ cells/mL. 100 μL 20% PEG8000 was added to 400 μL cell suspension. Then the cell suspension was vortexed for 15 s together with 400 μg of glass beads (Sigma, diameter 425–600 μm) and 3-10 μg of circular plasmid DNA. 100 μL of this cell suspension was spread onto a 1.5% agar plate (TAP medium with hygromycin B at 10 μg/mL). The plates were incubated for 10-15 days at 23 °C under 50 μmol photon m^−2^ s^−1^ light. Monoclones were then selected from surviving colonies and their foreign genes were verified, which were called strain PC and PCHP containing plasmid pHyg3-*cpcBA* and pHyg3-*cpcBA*-*ho*-*pcyA,* respectively.

### Southern blot analysis

*C. reinhardtii* was grown to mid-log phase (3–5 × 10^6^ cells/mL) in TAP medium and harvested by centrifugation at 3000 ×g for 10 min. Then the cells were resuspended with Buffer HS I of the Plant Genomic DNA Extraction Kit (TaKaRa). The resuspension was vortexed for 3 min after adding appropriate amount of glass beads (Sigma, diameter 425–600 μm). Then the gemoetic DNA of *C. reinhardtii* was extracted with the Plant Genomic DNA Extraction Kit according to the manufacturer’s instructions. Genomic DNA (5 μg) of the recipient strain PC and PCHP transformed with plasmid pHyg3-*cpcBA* and pHyg3-*cpcBA*-*ho*-*pcyA*, respectively, was digested with both of these two group restriction enzymes *Apa*I/ *Sal*I and *EcoR*I/ *Sac*I for 3 h. Then electrophorese was performed on a 1% agarose gel in 1 × TAE buffer at 120 V for 30 min. The DNA fragments were then transferred onto a nylon membrane (Solarbio) for 24 h. The southern blot analysis was completed with the DIG High Prime DNA Labeling and Detection Starter Kit I (Roche) according to the manufacturer’s instructions. *CpcBA*, *ho* and *pcyA* were amplified with high-fidelity DNA polymerase GXL (TaKaRa) and used for Dig-labeled probes after purification. Primers used in this study are listed in Table 1.

### quantitative real-time PCR (qRT-PCR) analysis

To isolate RNA, 3 × 10^7^ cells of *C. reinhardtii* at mid-log phase were harvested by centrifugation at 3000 ×g for 10 min. Total RNA was extracted with an E.Z.N.A.™ Yeast RNA Kit according to the manufacturer’s instructions (Omega). The quality of RNA was examined by gel electrophoresis on a 1% agarose gel. Then RNA was reverse transcribed to cDNA with a PrimeScript™ RT Master Mix (TaKaRa). Primers for phycocyanin (*cpcBA*), *ho*, *pcyA* from *Arthrospira platensis* FACHB 314 and *RACK* 1 from *C. reinhardtii* were designed for qRT-PCR analysis. And the *RACK* I was regarded as the housekeeping control gene. TB GREEN™ Premix Taq™ II (TaKaRa), cDNA and primers were used to prepare the reaction solution of qRT-PCR. Real time qPCR was performed on the realtime PCR detection system (lineGENE 9640, Bioer) using SYBR Green as a fluorescent dye. The reaction procedure was as follows: 95 °C for 30 s, followed by 40 cycles of 95 °C for 5 s and 60 °C for 30 s. Three biological replicates were made for each sample. Primers used in this study are listed in Table 1.

### Determination of chlorophyll content

The chl concentration of *C. reinhardtii* culture was determined with 80% acetone according to Porra’s formula (Porra et al., 1989). OD_647_, OD_664_ and OD_750_ were measured by a spectrophotometer to plot a curve.

### Low-temperature fluorescence spectrophotometry

Cells were harvested by centrifugation at 3000 ×g for 10 min. According to Akio’s method, the cells were then resuspended with fresh TAP medium containing 15% PEG4000 (Murakami, 1997). Samples were adapted in the dark for 15 min before measurement of the spectra. Resuspension was frozen in a liquid nitrogen Dewar vessel. Then it was loaded into the F-4600 fluorescence spectrophotometer (Hitachi, Japan). Data of emission spectra at 77K from 600 nm to 800 nm with a slit width of 5 nm and exciting wavelength of 435 nm (for Chlorophyll a) and 580 nm (for phycocyanin) were measured, at a total Chl concentration of 50 μg/mL.

### Measurements of photosynthetic parameters

The fluorescence and P700 parameters of PSII and PSI were measured by a Dual-PAM-100 measuring system (Heinz Walz GmbH, Effeltrich, Germany) (Zivcak et al., 2015). Cells were collected by centrifugation at 3000 ×g for 10 min. The pellet obtained was suspended and homogenized in fresh TAP medium to a Chl concentration of 50 μg Chl/mL. The samples were dark adapted for 10 min in a dark environment prior to measurements. The light curve was triggered by light intensities of 16, 39, 71, 132, 204, 286, 408, 578, 802, 1087, 1434 and 1857 μmol photon m^−2^ s^−1^ with 30 s at each light intensity. The transgenic strain PC, PCHP and the control strain cc849 taken from the 2^nd^, 4^th^, 6^th^, and 8^th^ day of low and high light were used for measurement. Moreover, there were two completely parallel biological repeats for each strain.

### Growth and biomass accumulation

Rate of growth and biomass accumulation of *C. reinhardtii* culture was measured under two experimental situations. The cells were cultured mixotrophically in TAP medium with illuminating by 25 (low-light) or 50 (high-light) μmol photon m^−2^ s^−1^ with a 12:12 h dark/light cycle and shaking at 100 rpm daily, respectively. Three samples including cc849, PC and PCHP were measured in each experimental group. And there were three parallel biological replicates for each tested line.

The transgenic alga lines to be used were activated in fresh medium with hygromycin B at 10 μg/mL in advance. Then the fresh culture was inoculated into 300 mL TAP medium without hygromycin B. The initial cell concentration was 6 × 10^5^ cells/mL. The absorbance of all three tested lines at 750 nm was measured by a Spectrophotometer daily to plot a growth curve in this 10-day experiment. Cells were collected by centrifugation at 3800 × g for 15 min on the 10^th^ day and washed twice with double distilled water. The dry weight was determined after drying overnight in a freeze dryer.

## Acknowledgments

We are grateful to Dr. Xiaonan Zang for her guidance on the progress of this work and the revision of the paper.

---

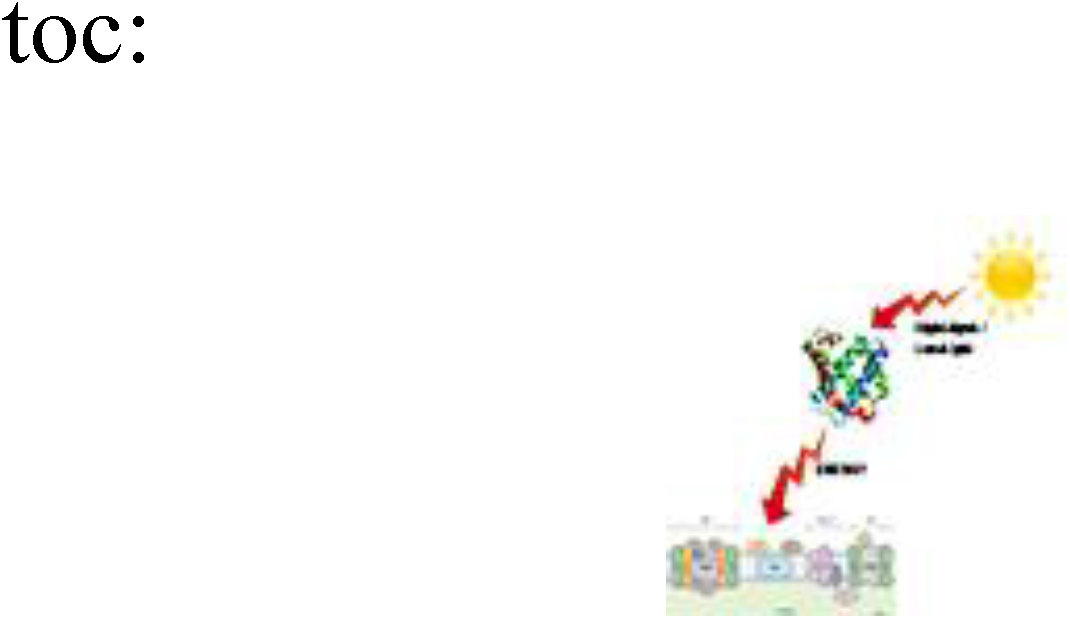

## References

Berthold P, Schmitt R, Mages WJP (2002) An Engineered Streptomyces hygroscopicus aph 7” Gene Mediates Dominant Resistance against Hygromycin B in Chlamydomonas reinhardtii. 153: 401–412

Biswas A, Vasquez YM, Dragomani TM, Kronfel ML, Williams SR, Alvey RM, Bryant DA, Schluchter WM (2010) Biosynthesis of Cyanobacterial Phycobiliproteins in *Escherichia coli*: Chromophorylation Efficiency and Specificity of All Bilin Lyases from *Synechococcus sp* Strain PCC 7002. Applied and Environmental Microbiology 76: 2729–2739

Cardol P, Gloire G, Havaux M, Remacle C, Matagne R, Franck F (2003) Photosynthesis and state transitions in mitochondrial mutants of *Chlamydomonas reinhardtii* affected in respiration. Plant Physiology 133: 2010–2020

Chapter 8 - Chlamydomonas in the Laboratory (2009). In EH Harris, DB Stern, GB Witman, eds, The Chlamydomonas Sourcebook (Second Edition). Academic Press, London, pp 241–302

Dejtisakdi W, Miller SM (2016) Overexpression of Calvin cycle enzyme fructose 1,6-bisphosphatase in *Chlamydomonas reinhardtii* has a detrimental effect on growth. Algal Research-Biomass Biofuels and Bioproducts 14: 116–126

Doi M, Shima S, Egashira T, Nakamura K, Okayama S (1997) New bile pigment excreted by a *Chlamydomonas reinhardtii* mutant: A possible breakdown catabolite of chlorophyll a. Journal of Plant Physiology 150: 504–508

Frankenberg N, Mukougawa K, Kohchi T, Lagarias JC (2001) Functional genomic analysis of the HY2 family of ferredoxin-dependent bilin reductases from oxygenic photosynthetic organisms. Plant Cell 13: 965–978

Franklin SE, Mayfield SP (2004) Prospects for molecular farming in the green alga *Chlamydomonas reinhardtii*. Current Opinion in Plant Biology 7: 159–165

Glazer AN (1989) Light guides. Directional energy transfer in a photosynthetic antenna. J Biol Chem 264: 1–4

Green BR, Parson WWJAiP, Respiration(2003) Light-Harvesting Antennas in Photosynthesis. 13

Griesbeck C, Kobl I, Heitzer M (2006) Chlamydomonas reinhardtii. Molecular Biotechnology 34: 213–223

Harris EH (2001) *Chlamydomonas* as a model organism. Annual Review of Plant Physiology and Plant Molecular Biology 52: 363–406

Hwangbo K, Ahn JW, Lim JM, Park YI, Liu JR, Jeong WJ (2014) Overexpression of stearoyl-ACP desaturase enhances accumulations of oleic acid in the green alga Chlamydomonas reinhardtii Plant Biotechnology Reports 8: 135–142

Johanningmeier U, Fischer D (2010) Perspective for the Use of Genetic Transformants in Order to Enhance the Synthesis of the Desired Metabolites: Engineering Chloroplasts of Microalgae for the Production of Bioactive Compounds. Bio-Farms for Nutraceuticals: Functional Food and Safety Control by Biosensors 698: 144–151

Kindle KL (1990) High-Frequency Nuclear Transformation of *Chlamydomonas-Reinhardtii*. Proceedings of the National Academy of Sciences of the United States of America 87: 1228–1232

Kothari R, Prasad R, Kumar V, Singh DP (2013) Production of biodiesel from microalgae *Chlamydomonas polypyrenoideum* grown on dairy industry wastewater. Bioresource Technology 144: 499–503

Kumar S, Hahn FM, Baidoo E, Kahlon TS, Wood DF, McMahan CM, Cornish K, Keasling JD, Daniell H, Whalen MC (2012) Remodeling the isoprenoid pathway in tobacco by expressing the cytoplasmic mevalonate pathway in chloroplasts. Metabolic Engineering 14: 19–28

Lapaille M, Thiry M, Perez E, Gonzalez-Halphen D, Remacle C, Cardol P (2010) Loss of mitochondrial ATP synthase subunit beta (Atp2) alters mitochondrial and chloroplastic function and morphology in *Chlamydomonas*. Biochimica Et Biophysica Acta-Bioenergetics 1797: 1533–1539

Lauceri R, Zittelli GC, Maserti B, Torzillo G (2018) Purification of phycocyanin from *Arthrospira platensis* by hydrophobic interaction membrane chromatography. Algal Research-Biomass Biofuels and Bioproducts 35: 333–340

Liu J, Zhang X, Sui Z, Zhang X, Mao YJJoAP (2005) Cloning and characterization of c-phycocyanin operon from the cyanobacterium Arthrospira platensis FACHB341. 17: 181–185

Lu Y, Zhang XJEJoB (2005) The upstream sequence of the phycocyanin β subunit gene from Arthrospira platensis regulates expression of gfp gene in response to light intensity. 62: 272–286

Murakami A (1997) Quantitative analysis of 77K fluorescence emission spectra in Synechocystis sp. PCC 6714 and *Chlamydomonas reinhardtii* with variable PS I/PS II stoichiometries. Photosynthesis Research 53: 141–148

Pittman JK, Dean AP, Osundeko O (2011) The potential of sustainable algal biofuel production using wastewater resources. Bioresource Technology 102: 17–25

Porra RJ, Thompson WA, Kriedemann PEJBba (1989) Determination of accurate extinction coefficients and simultaneous equations for assaying chlorophylls a and b extracted with four different solvents: verification of the concentration of chlorophyll standards by atomic absorption spectroscopy. 975: 384–394

Seo S, Okamoto M, Iwai T, Iwano M, Fukui K, Isogai A, Nakajima N, Ohashi Y (2000) Reduced levels of chloroplast FtsH protein in tobacco mosaic virus-infected tobacco leaves accelerate the hypersensitive reaction. Plant Cell 12: 917–932

Sidler WA (1994) Phycobilisome and Phycobiliprotein Structures. In DA Bryant, ed, The Molecular Biology of Cyanobacteria. Springer Netherlands, Dordrecht, pp 139–216

Su ZL, He DM, Qian KX, Zhao FQ, Meng CX, Qin S (2006) The recombination and expression of the allophycocyanin beta subunit gene in the chloroplast of *Chlamydomonas reinhardtii*. World Journal of Microbiology & Biotechnology 22: 101–103

Su ZL, Qian KX, Tan CP, Meng CX, Qin S (2005) Recombination and heterologous expression of allophycocyanin gene in the chloroplast of *Chlamydomonas reinhardtii*. Acta Biochimica Et Biophysica Sinica 37: 709–712

Sun DG, Zang XN, Guo YL, Xiao DF, Cao XX, Liu Z, Zhang F, Jin YM, Shi JW, Wang ZD, Li R, Yangzong ZX (2019) Cloning of *pcB* and *pcA* Gene from *Gracilariopsis lemaneiformis* and Expression of a Fluorescent Phycocyanin in Heterologous Host. Genes 10

Tan KWM, Lee YK (2017) Expression of the heterologous Dunaliella tertiolecta fatty acyl-ACP thioesterase leads to increased lipid production in *Chlamydomonas reinhardtii*. Journal of Biotechnology 247: 60–67

Tang K, Zeng XL, Yang Y, Wang ZB, Wu XJ, Zhou M, Noy D, Scheer H, Zhao KH (2012) A minimal phycobilisome: Fusion and chromophorylation of the truncated core-membrane linker and phycocyanin. Biochimica Et Biophysica Acta-Bioenergetics 1817: 1030–1036

Tetali SD, Mitra M, Melis A (2007) Development of the light-harvesting chlorophyll antenna in the green alga *Chlamydomonas reinhardtii* is regulated by the novel Tla1 gene. Planta 225: 813–829

Tooley AJ, Cai YPA, Glazer AN (2001) Biosynthesis of a fluorescent cyanobacterial C-phycocyanin holo-alpha subunit in a heterologous host. Proceedings of the National Academy of Sciences of the United States of America 98: 10560–10565

Wang F, Qi YF, Malnoe A, Choquet Y, Wollman FA, de Vitry C (2017) The High Light Response and Redox Control of Thylakoid FtsH Protease in *Chlamydomonas reinhardtii*. Molecular Plant 10: 99–114

Willows RD (2019) Chapter Five - The Mg branch of chlorophyll synthesis: Biosynthesis of chlorophyll a from protoporphyrin IX. In B Grimm, ed, Advances in Botanical Research, Vol 90. Academic Press, pp 141–182

Wlodarczyk LM, Dinc E, Croce R, Dekker JP (2016) Excitation energy transfer in *Chlamydomonas reinhardtii* deficient in the PSI core or the PSII core under conditions mimicking state transitions. Biochimica Et Biophysica Acta-Bioenergetics 1857: 625–633

Wollman FA (2001) State transitions reveal the dynamics and flexibility of the photosynthetic apparatus. Embo Journal 20: 3623–3630

Zhang J, Ma JF, Liu DS, Qin S, Sun S, Zhao JD, Sui SF (2017) Structure of phycobilisome from the red alga *Griffithsia pacifica*. Nature 551: 57–+

Zhang JL, Kang Z, Chen J, Du GC (2015) Optimization of the heme biosynthesis pathway for the production of 5-aminolevulinic acid in *Escherichia coli*. Scientific Reports 5

Zhou W, Sui ZH, Wang JG, Hu YY, Kang KH, Hong HR, Niaz Z, Wei HH, Du QW, Peng C, Mi P, Que Z (2016) Effects of sodium bicarbonate concentration on growth, photosynthesis, and carbonic anhydrase activity of macroalgae *Gracilariopsis lemaneiformis, Gracilaria vermiculophylla, and Gracilaria chouae* (*Gracilariales, Rhodophyta*). Photosynthesis Research 128: 259–270

Zivcak M, Brestic M, Kunderlikova K, Sytar O, Allakhverdiev SI (2015) Repetitive light pulse-induced photoinhibition of photosystem I severely affects CO2 assimilation and photoprotection in wheat leaves. Photosynthesis Research 126: 449–463

